# Biophysical boundaries for dormancy release in sunflower achenes

**DOI:** 10.1101/2024.12.28.630629

**Authors:** Gonzalo Joaquín Arata, Diego Batlla, Mailén Riveira-Rubin, Patricia Verónica Demkura, María Florencia Mazzobre, Guido Rolandelli, María del Pilar Buera, María Verónica Rodríguez

**Affiliations:** IFEVA, CONICET/Facultad de Agronomía de la Universidad de Buenos Aires, Av. San Martín 4453 (C1417DSE), Ciudad de Buenos Aires, Argentina; Cátedra de Cultivos Industriales, Departamento de Producción Vegetal, Facultad de Agronomía de la Universidad de Buenos Aires, Av. San Martín 4453 (C1417DSE), Ciudad de Buenos Aires, Argentina; Cátedra de Cerealicultura, Departamento de Producción Vegetal, Facultad de Agronomía de la Universidad de Buenos Aires, Av. San Martín 4453 (C1417DSE), Ciudad de Buenos Aires, Argentina; Cátedra de Fisiología Vegetal, Departamento de Biología Aplicada y Alimentos, Facultad de Agronomía de la Universidad de Buenos Aires, Av. San Martín 4453 (C1417DSE), Ciudad de Buenos Aires, Argentina; Universidad de Buenos Aires, Facultad de Ciencias Exactas y Naturales. Intendente Güiraldes 2160, Ciudad Universitaria, C1428EGA, Buenos Aires, Argentina; CONICET – Universidad de Buenos Aires, Instituto de Tecnología de Alimentos y Procesos Químicos (ITAPROQ). Intendente Güiraldes 2160, Ciudad Universitaria, C1428EGA, Buenos Aires, Argentina

**Author notes:** Corresponding authors: M.V.R. and G.J.A., María Verónica Rodríguez, Gonzalo J. Arata. Diego Batlla, Mailén Riveira-Rubin, Patricia V. Demkura, María del Pilar Buera, María Florencia Mazzobre, Guido Rolandelli.

**Keywords:** Sunflower Achene, Dormancy Release, Dry After-Ripening, Storage Temperature, Seed Moisture Content, Germination, Ageing, Glass Transition, Abscisic Acid

## Abstract

Reactions leading to dormancy release (DR) in “dry”, orthodox seeds are still poorly understood, as well as their dependence on moisture content and temperature. Sunflower achenes were used to explore the effects of MC combined with a wide range of storage temperatures (ST°) on DR dynamics, tested at 10 and 25°C. Embryo sensitivity to abscisic acid, oxygen uptake and ageing indicators were followed and complemented with predicted viability loss dynamics. Water status was addressed with sorption isotherms and the glass transition temperature (*Tg*). Small variations in MC and ST° had profound effects on achene dormancy and embryo responsiveness to ABA. Using a ‘temperature by humidity’ coordinate system we mapped three regions allowing DR within the glassy, rubbery and fluid states. Full DR (allowing germination at 10 and 25°C) occurred only in the rubbery state (above *T_g_*), and optimal ST° for full DR dropped from +30 to-18°C with MC increasing from 0.045 to 0.1 gH2O.gDW^-1^. Differently, under both glassy and fluid states, DR was incomplete (allowing germination at 25°C but not at 10°C) and required warm ST° which also promoted ageing.

**Highlight:** Sunflower achenes display diverse mechanisms for dormancy release in the “dry” state, depending on the physical state of water produced by different humidity and temperature combinations.

## Introduction

Sunflower (*Helianthus annuus* L.) is one of the main oilseeds worldwide, and the post-harvest conservation of sunflower achenes to be used in the next planting campaign is of outmost commercial importance. Seed dormancy is an internal block to germination with a clear adaptive value in wild species (Finch-Savage and Leubner-Metzger, 2006). However, in cultivated species prolonged dormancy can interfere with proper crop establishment (Benech-Arnold *et al*., 2012). This is a problem for sunflower hybrid “seed” (strictly, a fruit, achene or cypselae) which is usually dormant at harvest and requires a period of dry after-ripening to become non-dormant. This period may last from a few weeks to over a year depending on the genotype (Arata *et al*., 2021) and is influenced by the maternal (Bodrone *et al*., 2017; Lachabrouilli *et al*., 2021; Riveira Rubin *et al*., 2021) and post-harvest environments (Rodríguez *et al*., 2018; Bazin *et al*., 2011a). Dormancy attenuation during dry storage leads to a widening of the thermal range permissive for germination (Vegis, 1964; Batlla and Benech-Arnold, 2015) and non-dormant sunflower achenes can germinate at incubation temperatures between *ca* 10 and 30°C (Corbineau *et al*., 1990; Arata *et al*., 2021).

Orthodox seeds, like in sunflower, reach low moisture contents at dispersal/harvest maturity (Baskin and Baskin, 2004). In this low hydration range the cytoplasm can enter a glassy state at ambient temperatures. The capacity to form a glass is key for tolerating dehydration (Farrant and Hilhorst, 2021). Restricted molecular mobility in the glassy matrix is also essential to slow down reactions that cause ageing (Leopold *et al*., 1994; Ballesteros and Walters, 2011, 2019). Chemical reactions in the glassy matrix are limited to solid state oxidation, peroxidation, and carbonylation of molecules in proximity within the solid cytosol (Ballesteros and Walters, 2019). As these reactions lead to cumulative damage of seed components, seeds age and become non-viable. By increasing either temperature or moisture content (MC), the glassy matrix transitions to less viscous, “rubbery” and then “fluid” states, allowing higher molecular mobility including some enzyme reactions (Candotto Carniel *et al*., 2021). Because the effects of MC and storage temperature (ST°) on seed ageing dynamics are highly consistent across species, it was possible to develop the seed longevity model (Ellis and Roberts 1980). In contrast, the combined effects of MC and ST° on dormancy release (DR) are less clear when comparing studies across species, suggesting that reactions or mechanisms involved in dry after-ripening are more diverse than those involved in ageing.

Very often, and like ageing, DR is faster at higher temperatures (E.H. Roberts, 1965; Baskin and Baskin, 1976, 1986; Allen *et al*., 1995; R.J. Probert, 2000; Baldos *et al*., 2014). Besides, and different from aging, DR is favored within a particular MC range (corresponding to the second region of sorption isotherms) and is progressively inhibited with increasing hydration (Leopold *et al*., 1988; Esashi *et al*., 1993; Bair *et al*., 2006). In addition, several studies have reported interactions between MC and temperature on DR. For example, an inverse relationship between temperatures promoting DR and seed MC was observed in wild oat (M.E. Foley, 1994), while the opposite pattern was reported in *Arabidopsis* (Basbouss-Serhal *et al*., 2016) and sunflower (Bazin *et al*., 2011a). Nevertheless, these results may not be directly comparable due to different methodologies and ranges of MC and ST° explored in each study (Supplementary **Table S1**). A main concern about these and many other studies is that methodologies employed to manipulate MC are diverse and may affect the results and their interpretation. When allowing seeds to equilibrate with relative humidities (RH) generated by saturated salt solutions, changes in MC occur gradually over several days or even weeks, long enough to overlap temporally with significant changes in dormancy status. Another constraint is the narrow (or incomplete) range of storage conditions tested within the same study, leaving important interactions unnoticed. It is also unclear whether DR is favored by a particular state of water (glassy-rubbery-fluid). A better description of how MC and ST° influence dormancy release is needed to solve a practical agronomic problem (i.e., post-harvest conditions for sunflower hybrid seed production) but also to gain more insight into the mechanisms involved in the dry after-ripening process itself, in this and other species as well.

Whatever the changes that take place during dry storage, they have a strong impact on the physiology of the imbibed seed and its germination response (Chahtane *et al*. 2017). In Arabidopsis and barley, imbibed, dormant seeds exhibit differences in ABA metabolism and signaling as compared to imbibed, after-ripened (non-dormant) seeds (Ali-Rachedi *et al*., 2004; Benech-Arnold *et al*., 2006). In sunflower, ABA synthesis maintains embryo dormancy during achene development and maturation (Le Page-Degivry *et al*., 1990, 1996; Le Page-Degivry and Garello, 1992). However, ABA metabolism in mature, imbibed achenes was not clearly related with dormancy alleviation during after-ripening (Rodríguez *et al*., 2018). In contrast, sensitivity to ABA, assessed as the inhibition of embryo germination by exogenous ABA, decreased during dry storage (Bianco *et al*., 1994; Le Page-Degivry *et al*., 1996), and was associated with the different dormancy release dynamics under two contrasting storage temperatures (5 vs. 25°C; Rodriguez *et al*., 2018). The cause for this decreased responsiveness to ABA may involve direct changes in the functionality of components of the ABA signaling pathway, or other hormonal pathways known to antagonize ABA, such as gibberellins and ethylene (Corbineau *et al*., 1990, 2014; Corbineau and Côme, 2003; Xia *et al*., 2019).

The aims of this work were to investigate the effects of a broad range of ST° and MC combinations on dormancy release (and other related physiological traits) of sunflower achenes, and to associate the observed patterns with the physical state of water (as described by the moisture content isotherms and the glass transition temperature, *Tg*). Importantly, experiments were conducted in a way that manipulation of achene MC resembled practices in the seed industry (forced drying under air flow-non equilibrium conditions-) which also shortened the time to reach a target MC. This allowed us to track changes in dormancy status under fixed MCxST° conditions.

Using the state diagram obtained for the *Tg* as a coordinate system of ‘temperature by humidity,’ we mapped three regions within the glassy, rubbery and fluid states where DR occurred in a “full” or in an “incomplete” mode (i.e., allowing-or not-germination towards cooler temperatures). Our results support the existence of a specific mechanism for DR in the rubbery state operating at both ambient and sub-zero temperatures, where full DR largely anticipates ageing, in contrast with glassy or more fluid states where DR is promoted by warmer temperatures that also accelerate ageing.

## Materials and methods

### Plant material

Field trials were conducted in the experimental field at the Facultad de Agronomía de la Universidad de Buenos Aires (FAUBA) (34°35′37″S 58°29′03″O) during spring and summer in 2017-2018, 2018-2019 and 2019-2020 and are referred to as *experiments 1, 2* and *3*, respectively. Field plots were sown during early October, with plants in rows at a density of 5 plants/m^2^. Standard practices were followed for pest control, mineral nutrition and irrigation. The phenology of individual plants was followed according to Schneiter and Miller (1981). Experiments were performed with inbred line “600” (source: Instituto Nacional de Tecnología Agropecuaria, INTA), derived from Russian/USDA North Dakota, with oil content 49%. Dormancy phenotype of “600” is intermediate with no thermo-inhibition (Arata *et al*., 2021). At harvest maturity (moisture content *∼*11-12% fresh weight basis), 20-30 plants that did not differ by more than 5 d for R5.5 date were harvested. Heads were threshed manually, and achenes pooled for further analysis.

### Moisture content manipulation and storage treatments

Initial MC was determined gravimetrically. Each pool of achenes (>2.5 Kg) was then divided into four samples (*∼*500-600g) to obtain different target moisture content levels (MC ∼4, 6, 8, 10%, dry weight basis). Two different procedures were followed to manipulate MC (summarized in Supplementary **Fig. S1**). In *experiment 1,* four permeable polyamide bags containing 500 g achenes were placed in an experimental dryer (constant air flow at 35°C) to MC∼6%. Two of these samples with MC 6% were rehydrated to MC 8 and 10% by placing them in a saturated chamber (100% RH) for 10-21 d at 20°C. Increase in MC was monitored daily by weighting the samples. Another 500 g sample with MC 6% was further dried to MC 4% over silica gel (1:1 w/w, replenished with dry silica twice a day) at 20°C. In *experiments 2* and *3* (seasons 2019-2020, 2020-2021) a desorption sequence was followed immediately after harvest. Achenes in polyamide bags (∼500 g each, MC 12-13%) were placed in the experimental dryer (air flow at 35°C) and removed sequentially as each target MC was reached (10, 8 and 6%). Drying times ranged from 2 to 8 h for MC 8 and 6%, respectively. The lowest MC (∼4%) was obtained by further drying over silica for 3-4 d at 20°C.

Moisture content was determined gravimetrically as follows: Subsamples of 10-20 achenes were weighed before (fresh weight, FW) and after (dry weight, DW) oven drying at 130°C for 120 minutes (three replicates per sample or storage treatment). Moisture content is expressed as percent (%) relative to dry weight.

Storage treatments began immediately after completing MC manipulation. For each target MC, 10 g aliquots of achenes (∼150 achenes) were placed in glass containers (100 cm^3^ caramel bottles) and closed with rubber stoppers hermetically sealed with vacuum grease and tightened with parafilm. A factorial design (4 MC x 6-7 storage temperatures) was followed with triplicate containers for each MCxST° combination. Depending on the experiment and MC, ST° ranged from +5 to 25 or 30°C (with 5°C increments) and-18°C (freezer). Additional MC (3.5 and 5.2%) and storage temperatures were used in *experiment 3* (up to +50°C). Separate sets of containers were used for each sampling time (e.g. 30 and 70 d) in *experiments 1* and *2* so these remained closed until the germination assays at 30 and 70d. In *experiment 3* one set of containers was used, and these were opened and re-sealed repeatedly.

After *ca*. 200 d of storage, remaining achenes in *experiment 1* were used to test germination and seedling vigor (Supplementary **Fig. S3, S4** and **S5**). Achenes were incubated at 10°C until germination reached a plateau after 14 d. Achenes that did not germinate were transferred to Petri dishes with 6 ml of 100 µM ethephon (an ethylene donor) solution at 10°C, and germination was evaluated after four days. Germinated achenes were placed in plastic trays (10×15×4cm) over a cotton layer with filter paper soaked with distilled water and cultivated in a growth chamber (22-23°C, 16 h photoperiod) for four days. On day 4, seedlings were weighed individually. Data is presented for each storage treatment as the average FW of all seedlings ± SD (n = 20 to 25 seedlings).

### Storage under anoxia and oxygen measurements

Storage under anoxia was combined with MC 6 and 8% in *experiment 1* and with all MC levels in *experiment 2*. To generate anoxia, air was replaced by a constant flow of N_2_ gas through two (in/out) needles inserted through the rubber stopper for 1 min. After removing the needles, the outer surface of the rubber stopper was sealed with nail polish, and excess pressure was released before it solidified. These treatments were stored at +5 and 25°C. The oxygen level in the headspace of the glass containers with achenes was measured by inserting a needle-type micro-sensor through the rubber stopper (50 µm tips, Presens, Neurburg, Germany). All measurements were performed at 25°C (samples placed in a thermal bath). Data, obtained in mV, was converted to O_2_ concentration using a calibration curve (interpolated from O_2_ concentration in ambient air and anoxia). These values were then expressed as relative to ambient O_2_ concentration. Differences in O_2_ concentration during storage (measured at 30 and 70 d in *experiment 1*.) were used to calculate O_2_ uptake rate by achenes in each 100 cm^3^ container and expressed as umol O_2_ d^-1.^ 10 g of achenes^-1^.

### Germination tests

Dormancy level was assessed by incubating achenes and embryos at 10°C (which favors the expression of dormancy) and at 25°C (in *experiments 2* and *3*) or 30°C (*experiment 1*), where dormancy is less expressed (see Arata *et al*., 2021). Incubating at two contrasting temperatures allowed us to distinguish among deep-dormant samples (e.g., soon after harvest, where differences were best observed at 25°C) and among less-dormant samples where differences were best observed at 10°C. On each sampling date, and for each replicate container, germination assays were conducted with 25 achenes (or 20 embryos, after pericarp and seed coat removal) per Petri dish, placed on top of two layers of laboratory filter paper and 6 ml of distilled water and incubated for 15 d at 10 and 25°C (dark). Embryos were also incubated in 5 µM ABA (at 10°C in *experiment 1*, and 25°C in *experiment 2*). Germinated units were scored every 2-3 d and removed from the dish. Achenes were considered germinated when the radicle protruded 2-3 mm, and embryos when the radicle elongated >5 mm and began to curve. Germination data for each storage treatment is shown as the average of 3 replicate Petri dishes (from 3 storage containers) with the standard error of mean.

### Electrical conductivity of seeds

Twenty-five de-hulled seeds (pericarp removed, without damaging the seed coat) were placed in 37.5 ml of de-ionized water (< 2 μS.cm^-1^ g^-1^) at 25°C according to Szemruck *et al*. (2015). The electrical conductivity of the solution was measured after 24 h at 25°C using an Accumet AP85 portable conductometer, and results expressed in μS.cm^-1^ g^-1^.

### Quantification of endogenous ABA in embryo axes

Embryo axes were dissected from achenes that had been stored for 70 d (*experiment 2*), flash frozen in liquid N_2_ and stored at-80°C until processing for ABA quantification by radioimmunoassay (Steinbach *et al.,* 1995). Each replicate consisted of 20 axes from a storage container. Treatments included MC 4, 6, 8 and 10% in combination with ST° 5, 15 and 25°C. ABA is expressed as pg ABA.mg^-1^ dry weight of tissue.

### Moisture content shifting experiments

Freshly harvested achenes from *experiment 3* were dried to MC 6, 8 and 10% (experimental air dryer at 35°C) and stored at 20°C in hermetic containers (100 g achenes per MC). After 28 d of storage, germination was tested at 10 and 25°C, and 40 g sub-samples from each MC condition were used for MC shifting as follows: from 6 to 10% (11 d in humid chamber RH 100%), from 8 to 6% and from 10 to 6% (by drying under air flow at 35°C; 2-6 h). Once the new MC were reached, all samples continued storage in hermetic containers at 20°C. Lowering MC by drying under air flow was faster than rising MC in a humid chamber; therefore, achenes dried to MC 6% “after-ripened” for additional 9 d during gradual hydration of samples that reached MC 8 and 10%. Achene germination was tested immediately when MC shifting was complete for all treatments (39 d), and again at 60 and 100 d since the onset of storage.

Another trial was conducted to address the effect on DR of shifting MC at a constant temperature. Achenes with MC 4.2% stored 16 months at 10 and 15°C maintained a deep dormant state. For each ST°, 3 sub-samples were placed in a humid chamber (RH100%) until their MC reached 6%, and then stored hermetically for another 40 d. The corresponding ST° (i.e., 10 and 15°C) was maintained during MC modification and subsequent storage. Germination tests were conducted at 10 and 25°C.

### Sorption isotherms

Moisture adsorption isotherms were obtained for intact achenes and for embryo axes (native and defatted for *Tg* analysis). Samples were first dried for 1 d at 57°C under vacuum and then they were exposed to different relative humidities between 11 and 95%, using saturated salt solutions at 20°C in hermetic closed containers. After equilibrium, water activities (*a_w_*) of samples were measured using an electronic dew-point water activity meter (Aqualab Series 3, Decagon Devices, Pullman, WA). Final MC was determined after drying for 2 d at 57°C under vacuum. All systems were analyzed in triplicates. The isotherm parameters (monolayer, C, K) were estimated by fitting the Gugenheim-Anderson-de Boer (GAB) equation.

Sorption isotherms were obtained for achenes with different MC (4 – 11%) obtained using the same methodology as in the physiological experiments (i.e., achenes hydrated to MC14% were subject to forced drying at 35°C). Isotherms were obtained after measuring the relative humidity (RH) of air in equilibrium with achene samples in closed containers (40 g in 100 cm^3^), which were sequentially placed at different temperatures (-18 to 30°C) until RH stabilized.

### Glass transition temperature determination by DSC and ^1^H-NMR

Embryo axes were defatted in chloroform:methanol 2:1 for 24h to reduce signals from lipid transitions, as in Williams and Leopold (1989), and equilibrated over saturated salt solutions (HR 11-95%) at 20°C to obtain different MC (2.45 – 20.83%). Glass transition temperature (*Tg*) was determined by differential scanning calorimetry (DSC) and ^1^H-nuclear magnetic resonance (^1^H-NMR). A differential scanning calorimeter (Mettler Toledo, model 822, Zürich, Switzerland) was used to evaluate thermal transitions of embryo axes according to Farroni and Buera (2014). Briefly, about 10 mg of sample, exactly weighted, with different MC (%, dry basis) were hermetically sealed in aluminum pans and analyzed under nitrogen atmosphere in the range between-100°C to 120°C at a heating rate of 10°C/min. An empty aluminum pan was used as reference and samples were analyzed in duplicates. Thermograms were evaluated using STARe software v. 6.1 (Mettler Toledo, Zürich, Switzerland). Data is presented as *Tg* onset obtained for individual runs (2-4) for each MC sample.

Molecular mobility was analyzed by ^1^H-NMR, measuring the spin-spin transverse magnetization relaxation times (*T_2_*). A pulsed nuclear magnetic resonance instrument (Bruker, model Minispec mq 20, Karlsruhe, Germany) with a 0.47 T magnetic field and operating at a frequency of 20 MHz was used. Probe head was kept at 40 ± 1°C. Equilibrated samples were placed in 10 mm diameter glass tubes, tightly compressed to avoid air holes, and heated from 10°C to 80°C at 10°C intervals using a thermal bath (Haake, model Phoenix II C35P, Thermo Electron Corporation, Waltham, MA). Glass tubes were sealed to prevent moisture loss during heating. Free induction decay (FID) of protons was analyzed after a single 90° pulse of 2.74 µs, with an acquisition period of 0.5 ms, recycle delay of 3 s with 4 scans and acquiring a total of 250 data points. Signals from this sequence are related to protons populations from matrix solids and water that have strong interactions with solids and thus, present short relaxation times (typically lower than 30 microseconds). ^1^H-NMR signals were fitted to mono-exponential equation:

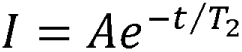

where *I* indicate the signal intensity at time *t*, *T_2_*is the relaxation time of protons and *A* is proportional to the number of protons associated to the signal. *T_2_*values (ms) were plotted against temperature to analyze matrix mobility and the related glass transition temperature (*T_g_*) (Farroni *et al*., 2008).

### Estimation of viability

The improved viability function of Ellis and Roberts (1980) was used to calculate achene viability during storage at each MC and ST° condition. The viability equation is:

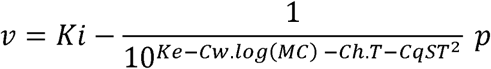

where *v* is the viability (in probits) after *p* storage days of storage, *Ki* the initial germination (probits), *MC* the moisture content (% FW basis), *T* the storage temperature (°C) and *Ke*, *Cw, Ch* and *Cq* species-specific constants (6.74, 4.16, 0.0329, 0.000478 respectively). A *Ki* value of 2.3 probits was selected for calculations, being this value in the upper range reported for a variety of commercial sunflowers (Saux *et al*., 2020). Dormancy release dynamics were obtained by adjusting non-linear functions (Gaussian or Gompertz) to final germination (%) data at different storage times (from 0 to ∼100 d). Storage time to reach 50% germination (T50) was obtained from the adjusted functions and used to calculate the DR rate (DRR = 1/T50). Values for DRR were expressed as relative to the maximum value observed at each incubation temperature (either 10 or 25°C).

### Data analysis

First processing of all data (including germination and viability percentages, oxygen levels and uptake rates, EC, ABA content, and moisture content values) was performed in Excel. Fitting of non-linear functions (for DR and viability dynamics, sorption isotherms, and *Tg*), ANOVA and multiple comparisons (for germination data, ABA levels), and data presentation was performed with GraphPad Prism 7 software (H.G. Motulsky, 2003).

## Results

### Effect of pre-storage manipulation on initial dormancy level

The dormancy status of achenes was assessed immediately after adjusting their MC and prior to storage. Achene drying partially reduced embryo dormancy, as shown in *experiment 2* (Supplementary **Table S2**). Achene germination was 16% for MC 4.7% and remained null for the other MC levels, while germination of embryos in water at 25°C reached 65, 36, 3.3 and 3.3% for MC 4.7, 6.6, 7.9 and 10%, respectively. When tested at 10°C (a condition that enhances expression of dormancy in sunflower), pre-storage germination scores were equally null for achenes and embryos from all MC levels. Besides this mild attenuation of dormancy by drying to MC 4 and 6% (detected at 25°C), a deep dormant state was still evident for all MC. This allowed us to relate any further changes in dormancy to subsequent storage conditions. Achene MC remained mostly unchanged during storage in the hermetic containers and was not significantly altered by ST° (Supplementary **Fig. S1**).

### Effects of MC and ST° on dormancy release

Dormancy release (DR), assessed through germination of achenes and embryos at 30 and 70 DAS, was deeply affected by ST° and MC with strong interactions between both factors (**Fig. 1**). For the lowest MC (4.7%, **Fig. 1A-D**), DR was only observed after storage at 25°C (i.e., higher achene germination values together with increased embryo germination in water and 5 uM ABA) while storage at 20°C and below maintained deep embryo dormancy. In contrast, DR was strongly promoted in achenes with MC 6.6% when stored between 5 and 20°C (resulting in high germination values at both 25 and 10°C incubation; Fig. **1E-H**). Nevertheless, also for MC 6.6%, DR was slower under storage at 25°C (resulting in lower germination values for achenes imbibed at 10°C and for embryos in 5 uM ABA; Fig.**1E-G**). This negative response to warmer ST° was more pronounced in achenes with MC 7.9% (Fig. **1I-L**) stored between 5 and 25°C. This pattern was also followed by isolated embryos incubated in water at 10°C (30 DAS; Fig. **1I**) and in 5 uM ABA at 25°C (Fig. **1K**).

**Figure 1.**
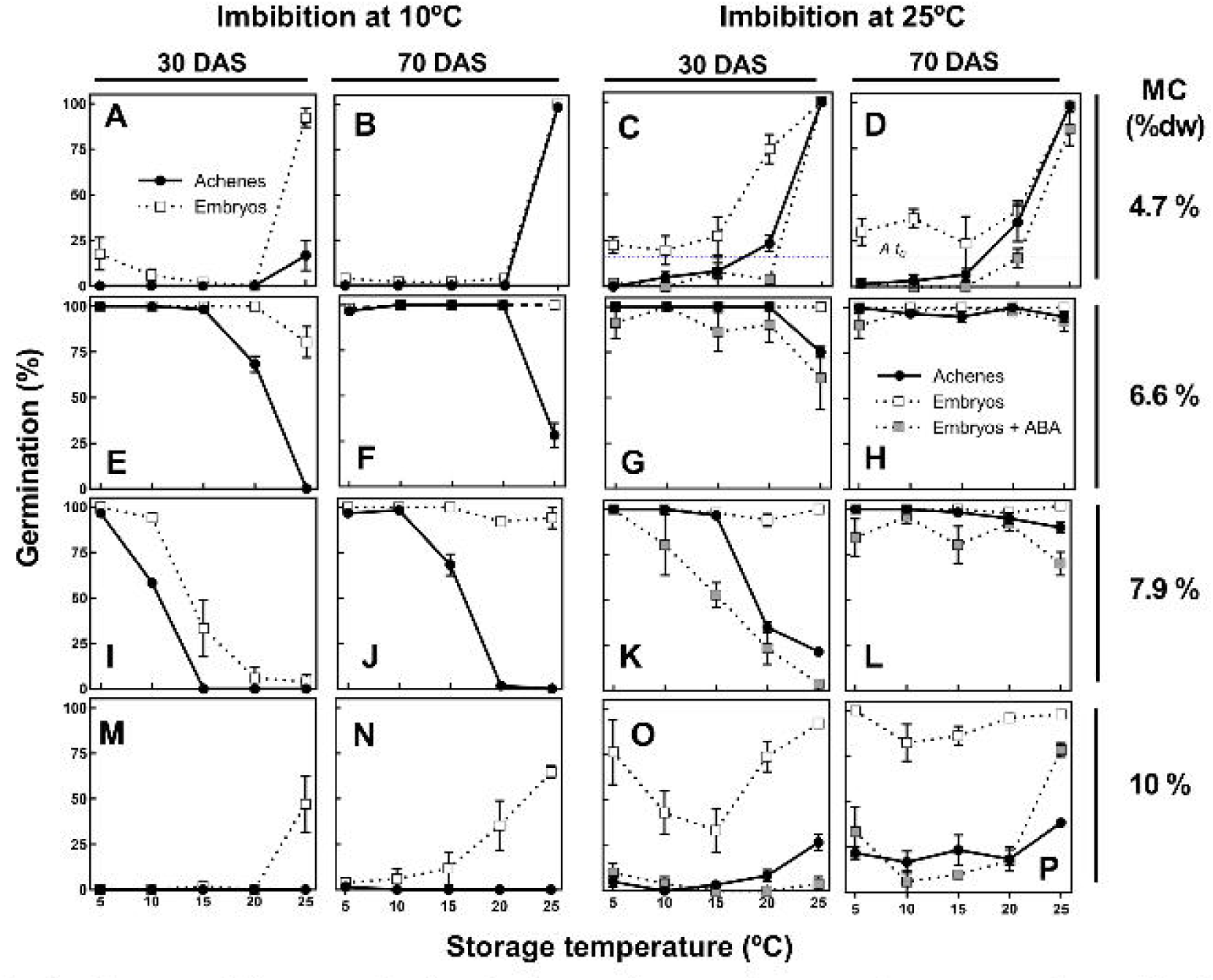
Effect of achene moisture content and storage temperature on dormancy release. Each panel shows final germination percentage of sunflower achenes and embryos as a function of ST° (+5, 10, 15, 20 and 25°C). Different achene moisture contents are presented in rows (from top to bottom, MC: 4.7, 6.6, 7.9 and 10.0% dry weight basis). Germination was tested at 10°C (left columns) and 25°C (right columns), at 30 and 70 days after storage (DAS). Achenes were incubated in water (filled circles), and embryos in water (empty squares) and in 5 µM ABA (grey squares, only at 25°C incubation). Pre-storage achene germination (*At_0_*) is shown with a horizontal dashed line (when line not visible, germination was null). Each data point represents the mean ± S.E.M. (n=3 replicate storage containers, each tested in one replicate dish with 25 achenes/embryos). Data belongs to *experiment 2*.

For MC 10%, embryo dormancy was partially alleviated by increasing ST>15°C, although achene germination values remained low at both incubation temperatures (**Fig. 1M-P**). Interestingly, a partial promotion of DR in response to increasing temperatures above 15°C was also observed in *experiment 1* for achenes with high MC (8-10%). In this case, achenes germinated when imbibed at 30°C, but failed to germinate at 10°C, even after prolonged (7 months) storage (Supplementary **Fig. S2** and **S3**). These results altogether support that high MC combined with warm ST (>15°C) lead to an incomplete alleviation of dormancy, allowing achene germination-if any-only at warm incubation temperatures (25-30°C) but not at 10°C.

### Achene dormancy correlated with embryo responsiveness to ABA

Achene germination was closely followed by embryo germination in 5 uM ABA (either when tested at 10 or 25°C; see also *experiment 1,* Supplementary **Fig. S2**). This led to a positive and significant correlation between germination scores of achenes in water and embryos in 5uM ABA (p<0.0001, r=0.96; **Fig. 2A**), including all combinations of MCxST and sampling times (30 and 70 DAS). These results support that changes in achene dormancy under different storage conditions are driven by changes in embryo sensitivity to endogenous ABA. The only deviation from this pattern was for MC10%-ST25°C at 70 DAS.

**Figure 2.**
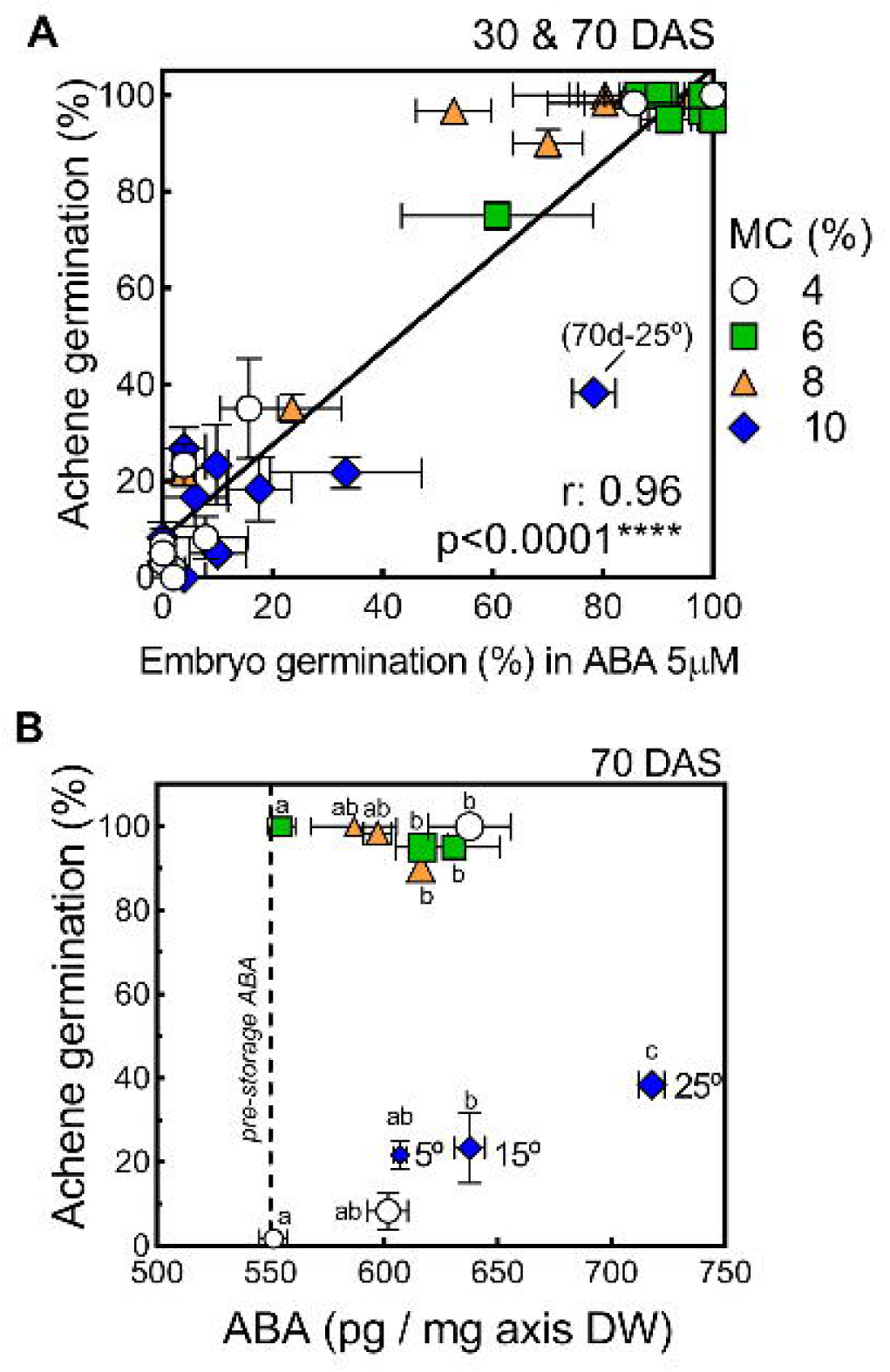
Achene germination correlated with embryo germination in ABA, but not with ABA content. (**A**) Biplot for achene germination in water and embryo germination in 5 µM ABA, both incubated at 25°C. Data points belong to different MCxST° treatments tested at 30 and 70 DAS (data from Fig. 1, *experiment 2*). Pearson’s correlation coefficient (r) and p-value are shown. Symbol shapes in **A)** and B) (circles, squares, triangles and diamonds) indicate moisture content (4.7, 6.6, 7.9 and 10.0% dry weight) (**B**) Achene germination % at 25°C in water versus ABA content in the embryo axis. Data belongs to achenes with different MC% (same as in **A**) stored at 5, 15 and 25°C, 70 DAS. Symbol size (larger) relates to ST° (higher). Vertical, dashed line indicates pre-storage ABA content. Multiple comparisons for ABA content data were performed by Tukeýs test; significant differences (p<0.05) among means are shown in different letters. Each data point in **A)** and **B)** is the mean ± S.E.M. of n=3 replicates (storage containers).

Endogenous ABA content in the embryo axis was also examined in all four MC levels after storage at 5, 15 and 25°C (**Fig. 2B**). After 70 DAS, ABA content had increased with ST° as compared to pre-storage levels (550 pg ABA.mg^-1^DW). This increase was highest (*ca* 30%) for achenes with MC10%-ST25°C (717 pg ABA.mg^-1^DW, p<0.0001). The observed changes in ABA content were not related to achene germination (p = 0.79, r =-0,043).

### Full dormancy release was not affected by oxygen availability

A possible effect of oxygen on DR under different MCxST° combinations was investigated by replacing air in the containers with gaseous N_2_ at the onset of storage (anoxia). The presence or absence of O_2_ had no impact on DR of achenes with MC∼4, 6, 8% stored at either 5 or 25°C for 70 d (and germination tested at 10 and 25°C; **Fig. 3**). In contrast, storage under anoxia improved achene germination for MC∼10% when tested at 25°C (germination at 10°C was null). This difference did not apply to embryo germination in water or in 5uM ABA, suggesting that the gaseous environment had no effect on embryo dormancy status. The lower germination of achenes with MC10% stored under normoxia as compared to anoxia may result from ageing and subsequent failure of the mechanism leading to pericarp opening, and which may involve a functional, living endosperm. This is consistent with increased electrical conductivity (indicative of damage in the endosperm/seed coat) tested 70 DAS in seeds from achenes with MC∼10% stored at 25 and 30°C for 70 d (Supplementary **Fig. S4B**). Reduced vigor caused by ageing might also explain the discrepancy between germination of achenes (in water) and embryos (in ABA) for MC10%-ST25°C tested 70 DAS, observed in **Fig. 2A**.

**Figure 3.**
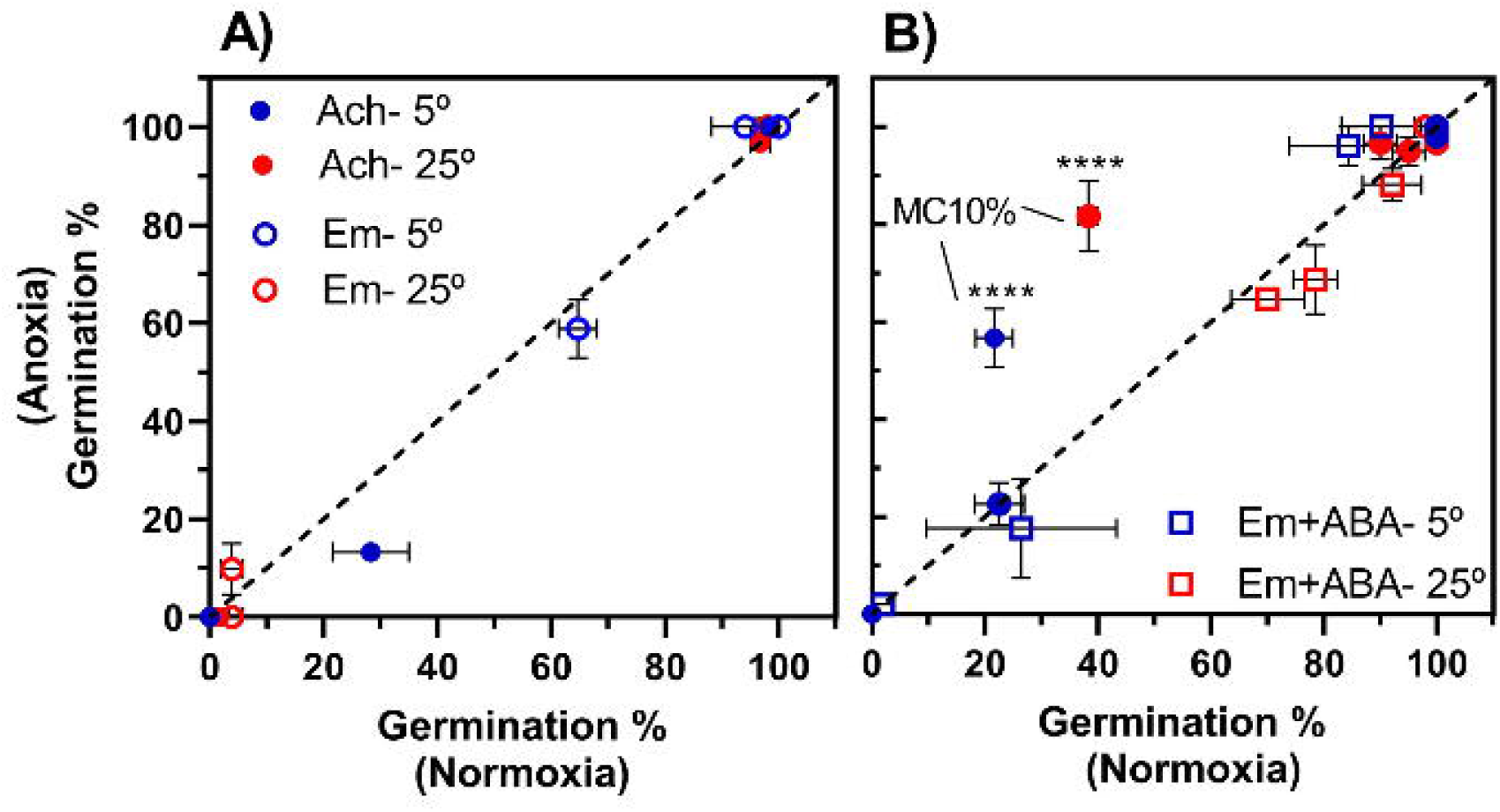
Effect of oxygen availability on dormancy release. Biplots for germination values obtained for achenes 70 DAS stored in closed containers with N_2_ (anoxia) or air (normoxia). Germination was tested at 10 **(A)** and 25°C **(B)**. Storage under anoxia/normoxia conditions were combined with four different MC (ca 4, 6, 8 and 10% dry weight) and two ST° (5 and 25°C; blue and red symbols). Final germination (%) is shown for achenes in water (Ach), embryos in water (Em) and in 5uM ABA (Em+ABA, only tested at 25°C). Each data point is the average of three replicate storage containers (mean ± S.E.M.). Data belongs to *experiment 2*. A significant difference (p<0.0001) between anoxia and normoxia was observed only for achenes with MC10% under each ST° when tested at 25°C (2-way ANOVA, Sidak‘s multiple comparisons test).

The O_2_ level in the headspace of closed containers was measured 70 DAS (**Fig. 4a**). The rates of O_2_ uptake are presented along an RH gradient (**Fig. 4b**) to allow comparisons with the moisture sorption isotherms shown below (Fig. **4c**). For achenes with MC 4-8% (spreading along region 2 of the sorption isotherms), O_2_ uptake rates were null or very low at ST° below 15°C but increased progressively (and similarly among MC) with ST°20-25°C. This led to a 10-15 % drop in O_2_ levels 70 DAS, which may be explained by spontaneous auto-oxidation of seed components. A similar decrease in O_2_ level was observed for MC10% and ST° up to 15°C (also within region 2), but O_2_ uptake rates increased sharply at higher ST (>15°C), and O_2_ was almost depleted 70 DAS at 30°C. Oxygen uptake under these conditions can be attributed to partial reactivation of respiration in the seed (and of microorganisms in the pericarp) consistent with RH>80% (**Fig. 4b**) and the beginning of region 3 of the sorption isotherms at 20-30°C (**Fig. 4c**).

**Figure 4.**
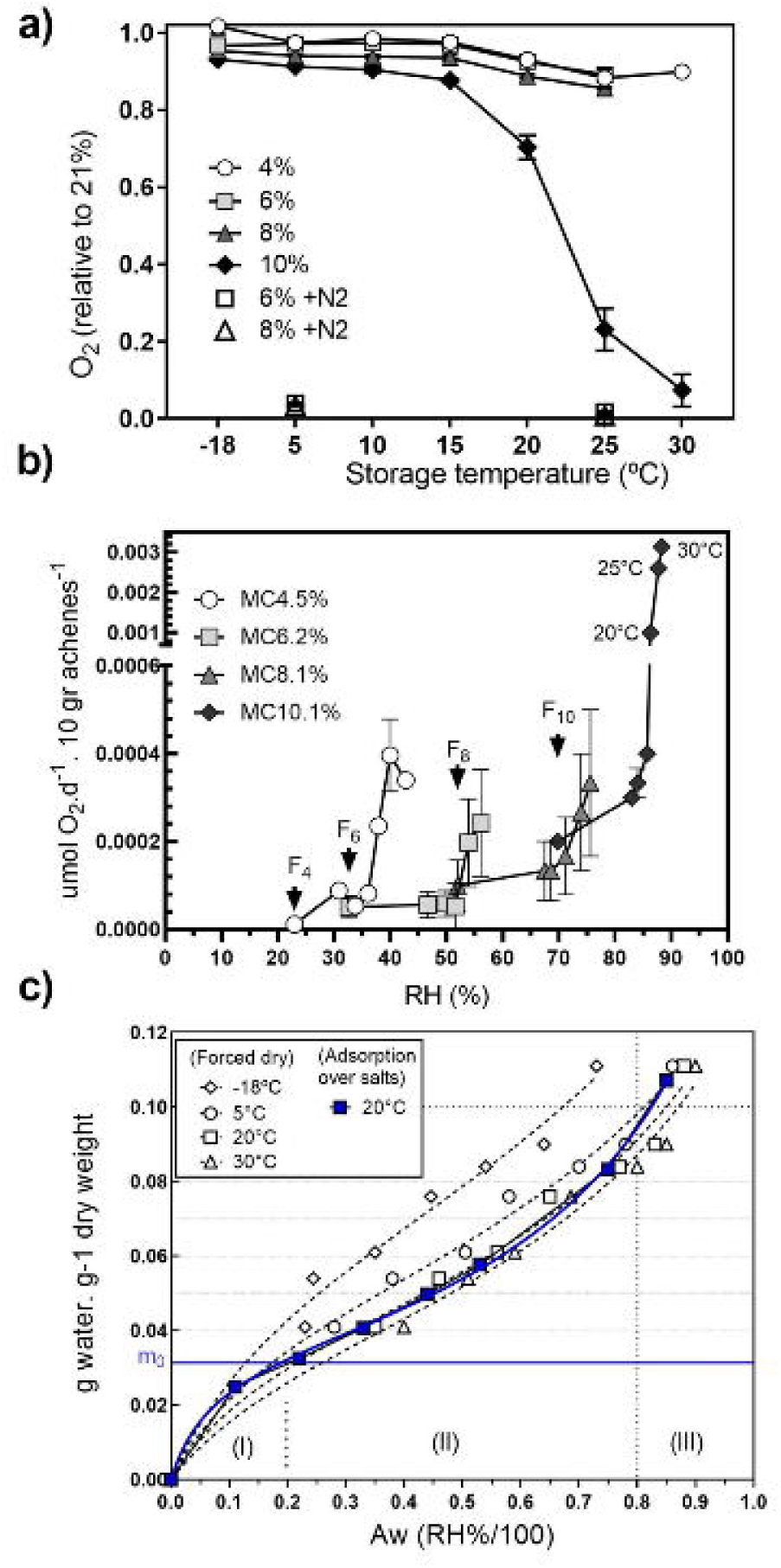
Oxygen uptake and RH% affected by achene MC and ST°. **(a)** Oxygen (relative to air) in the headspace of hermetic 100 cm^3^ flasks containing achenes (10 g) with different MC (ca 4.5, 6.2, 8.1 and 10.1%; filled circles, squares, triangles, diamonds) and stored at constant temperatures (-18 to 30°C) for 70 d. Each data point is the average ± SEM (n=3 replicate containers). Oxygen was also measured in containers initially flushed with N_2_ (anoxia treatments; empty triangles and squares). Data belongs to *experiment 1*. **(b)** Rate of O_2_ uptake as a function of RH (%) inferred from the sorption isotherms below. Arrows point to RH at-18°C (freezer, FMC) for each MC. For consecutive data points of a same MC series, ST° increases from left to right (ST° values 20-30°C are indicated for MC10%). **(c)** MC isotherms obtained by measuring the RH (%) in equilibrium with a fixed achene MC (obtained by forced drying) and equilibrated at different temperatures (-18, +5, 20 and 30°C). The adsorption isotherm for achenes equilibrated over saturated salt solutions at 20°C is shown in blue (filled squares). After fitting the GAB model the estimated monolayer value was 0.032 g water. g^-1^ dry weight (horizontal blue line).

### Dormancy release and viability dynamics at different MCxST combinations

A new experiment (*experiment 3*) was performed to track temporal changes in achene dormancy along a 100-d storage period for different MCxST combinations including sub-zero (-18°C) storage temperature (see also Supplementary **Fig. S6** for DR dynamics in *experiment 2*). **Fig. 5** presents final achene germination values tested at 10 and 25°C and the adjusted dynamics for four achene MC levels (4.2, 6.6, 8.0 and 10.6%) combined with ST-18, 5, 10, 15, 20 and 25°C. Full DR is understood as achenes being able to germinate at both 10 and 25°C incubation, in contrast to incomplete DR, where achene germination proceeds at 25-30°C but not at 10°C. Full DR only occurred under discrete MCxST° combinations, and the range of ST° allowing full DR depended on MC%. For MC 6.6%, the ST° range allowing full DR was the widest (ST 5-25°C), with an optimal at 10-20°C. Outside this range full DR still occurred but was delayed, either towards lower (5°C) or higher (25°C) ST°, or was completely inhibited at ST-18°C. For MC 8% full DR took place at ST 5-15°C with an optimal around 5°C, and incomplete DR occurred at ST 20-25°C. For MC10%, notably, full DR was only observed during storage at - 18°C and was inhibited at ST 5°C or above. At low MC (4.2%) incomplete DR occurred at ST 25°C. The effect of higher ST° (25-50°C) was also explored in achenes with MC 3.5% and 5.2% (Supplementary **Fig. S7**). When MC 3.5% the only ST° promoting incomplete DR was 50°C, which also caused rapid ageing. In contrast, for MC 5.2%, full DR was promoted by ST° between 20 and 40°C, with the optimal ST° around 30°C (**Fig. S7**).

**Figure 5.**
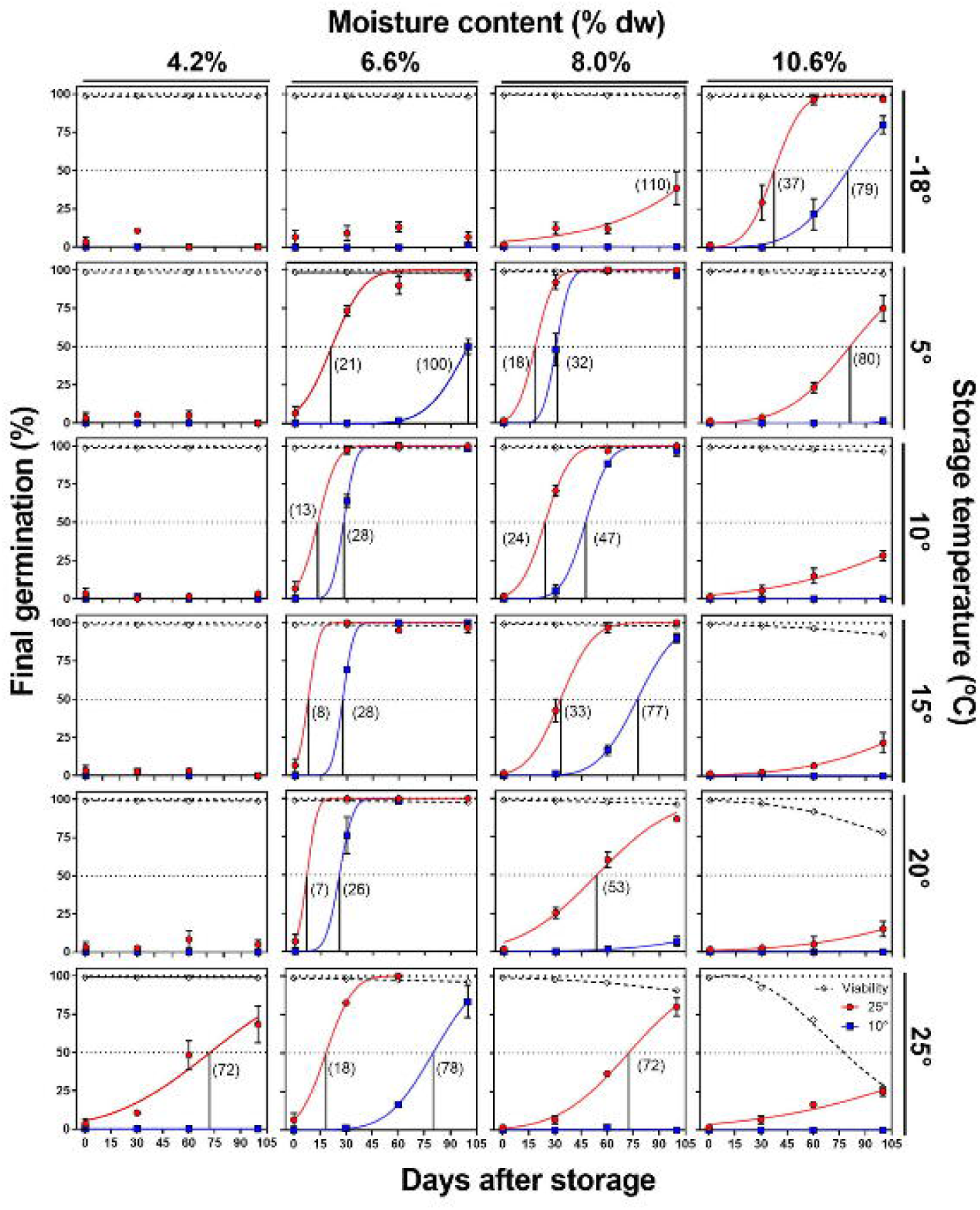
Dormancy release and predicted viability loss dynamics during storage at different MCxST° conditions. Each panel shows the final achene germination percentage at two imbibition temperatures (10 and 25°C, blue squares and red circles respectively) as a function of storage time (0, 30, 70 and 100 d) with their fitted non-linear functions. Each different panel presents data for a particular MC (in columns: 4.2, 6.6, 8.0 and 10.6% dry weight basis) and storage temperature (in rows:-18, +5, 10, 15, 20, 25°C). Data belongs to *experiment 3*. Storage time to reach 50% germination (T50) at either 10 or 25°C incubation, is shown in parenthesis. Similar dynamics were obtained for *experiments 1* and *2* (Fig. **S6**). Predicted viability (empty diamonds) were calculated for each storage condition using the viability function (Roberts and Ellis, 1980) with a Ki = 2.3. Each data point represents the mean ± S.E.M. (n=3 replicate containers).

The predicted viability curves were plotted together with DR dynamics for each MCxST° condition. Under conditions optimizing full DR (e.g., MC 6.6% at 10-15°C, or MC 8.0 at 5°C), DR largely anticipated ageing. In contrast, under storage conditions promoting incomplete DR at high temperatures (25°C and above), the DR dynamics overlapped more closely with ageing (e.g., MC 10% at 25°C, or MC 3.5% and ST 50°C).

### Optimal ST° for dormancy release was inversely related to achene MC

Based on the DR dynamics shown in **Fig. 5** and Supplementary **Fig. S7**, The DR rates (1/T50) were obtained and are shown in **Fig. 6**. Data is presented as relative to the maximum value observed in the whole dataset. For achenes with MC 3.5%, only partial DR (allowing germination at 25°C incubation) was observed after storage at 50°C. The minimum ST° allowing at least incomplete DR (leading to germination at 25°C but not at 10°C) shifted to 25°C for MC 4.2%. Full DR was promoted in achenes with MC 5.2% stored above 15°C, with an optimal at 30°C. Supra-optimal ST° (35-40°C) delayed DR (evident when tested at 10°C incubation). For MC 6%, DR was promoted by ST 5-25°C, with an optimal around 15-20°C. At higher MC (8 and 10%) ST° promoting DR were clearly shifted towards lower (even sub-zero) values. For MC 8%, the optimal ST° was ≤ 5°C, while for MC 10%, the optimal appeared to be well below zero.

**Figure 6.**
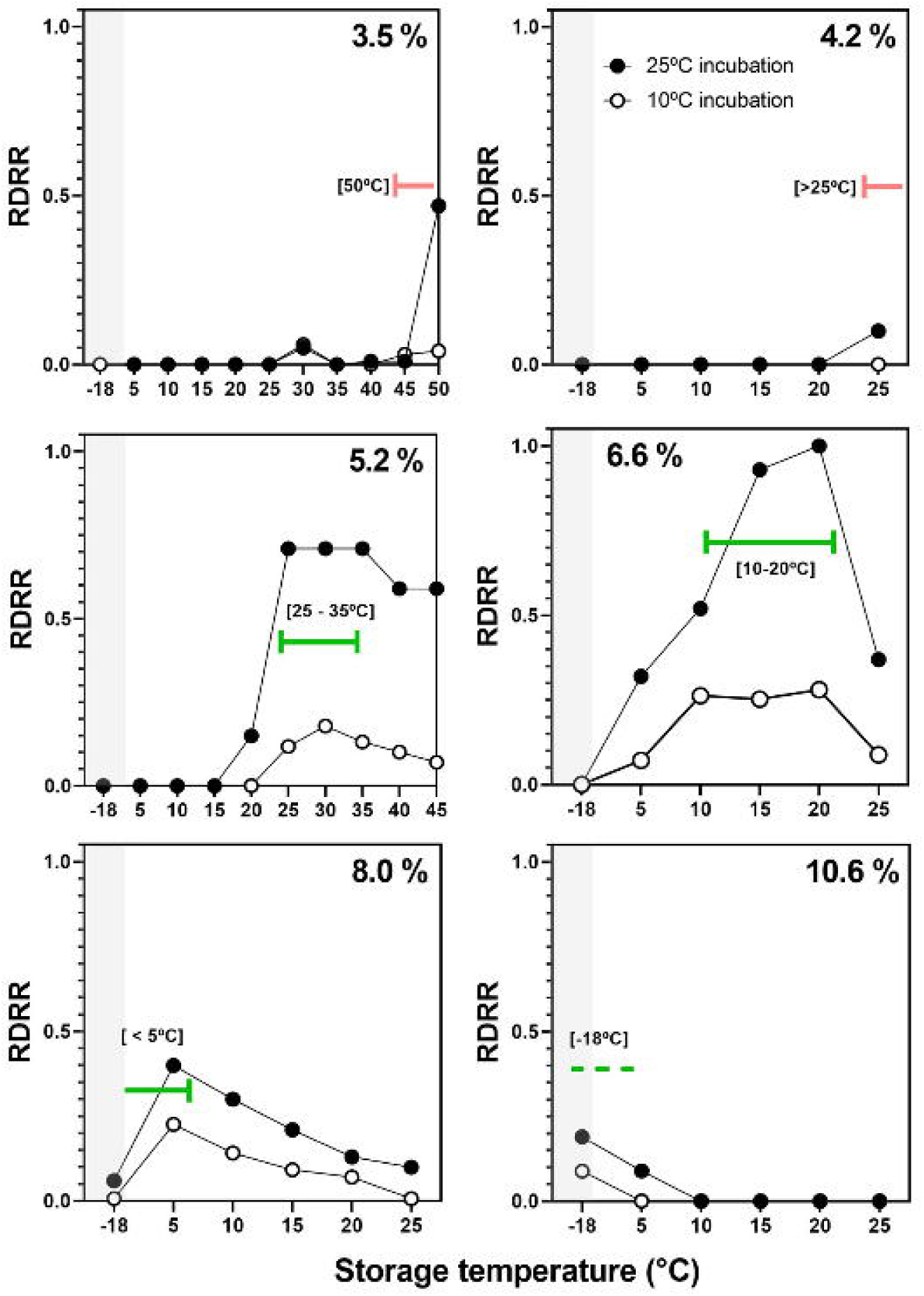
Relative dormancy release rate (RDRR) as a function of storage temperature for achenes with different MC. Relative DRR are presented for achenes with fixed MC values (3.5, 4.2, 5.2, 6.6, 8.0 and 10.6 % dry weight basis) and stored for up to 3 months at different temperatures (ST°) ranging from-18°C to +50°C. Germination was tested periodically at 10 and 25°C. Dormancy release rate values were calculated as 1/T50, and expressed as relative to the highest value (observed for MC6.6-ST20°C). The horizontal segment indicates the range of ST° promoting DR for each MC; green color indicates “full” DR, and pink indicates partial DR (germination occurs only at 25°C). All data belongs to the same achene pool (*experiment 3*).

### Optimal temperatures for dormancy release and the glass transition temperature (T_g_)

The optimal storage temperatures for DR were strongly dependent on achene MC and decreased from +30°C for MC *∼* 5% to about-18°C for MC *∼*10%. This inverse relationship between optimal ST° for DR and achene MC resembled a state diagram. To better understand if MC and ST° conditions promoting DR are related to a particular state of cellular water, the *T_g_* was measured in embryo axes. To avoid masking thermal transitions by signals from lipids, embryo axes were defatted before DSC and ^1^H-NMR analyses (Williams and Leopold, 1989). Sorption isotherms were used to relate MC of defatted axes with MC of native, whole achenes as used in DR assays (Supplementary **Fig. S8**). The isotherms also allowed us to estimate the monolayer MC (after fitting the GAB model; **Fig. 4c**). The obtained values for *Tg* are presented as a function of achene MC in **Fig. 7** together with the optimal temperatures for DR inferred from **Fig. 6**. A theoretical “longevity” line (Tmax_(v95%-90d)_) is also presented referring to the maximum ST° maintaining ∼95% viability for 3 months as a function of achene MC. **Fig. 7** shows that full DR takes place along a discrete range of MCxST conditions above the *T_g_*, within a non-glassy, “rubbery” state. Lower and upper limits can be set to the MC range allowing full DR. A minimum MC of ∼4.5% (MC_min_) was determined empirically (full DR was not observed below this MC value). This is also the MC at which the adjusted functions for T_opt (DR)_ and *T_g_* converge. At any MC below MC_min_, water is in a glassy state even at high temperatures (∼40-50°C). In the glassy state, ST° high enough to promote incomplete DR also promote rapid ageing (e.g., MC 3.5% and ST 50°C; Supplementary **Fig. S7**, **Table S3**). Increasing MC above MC_min_ leads to a rubbery state at “ambient” temperatures (below 35°C) but also at sub-zero temperatures, as observed by the *T_g_*dropping from +35°C to-80°C as MC rises from 4.5 to 10.5%.

**Figure 7.**
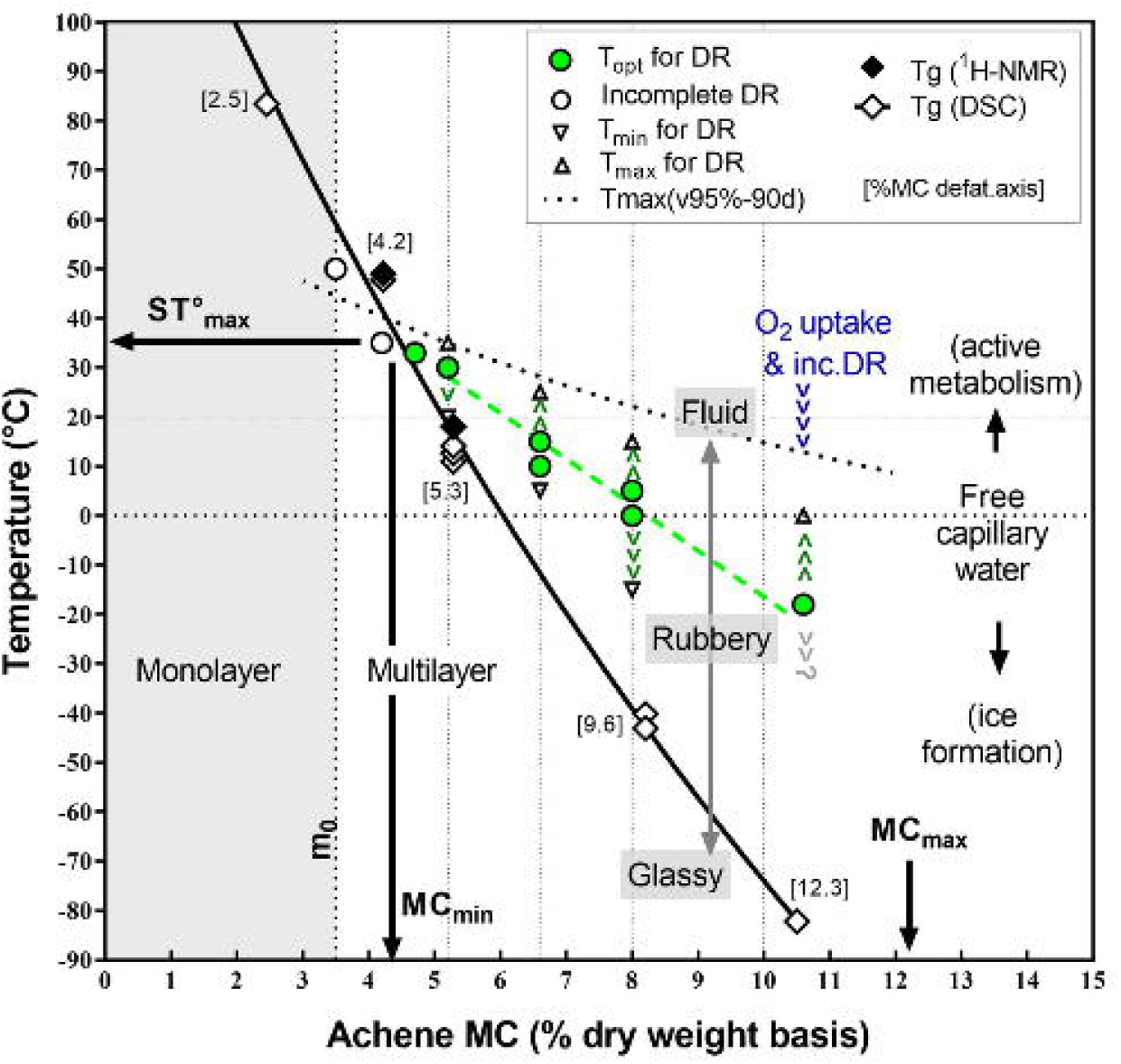
Cardinal temperatures for dormancy release and the glass transition temperature (*Tg*) as a function of achene MC. The *T_g_* was measured in embryo axes (by DSC-white diamonds-and ^1^H-NMR-black diamonds-) and the adjusted function is shown as a blue solid line along the achene MC (%, dry weight basis) axis. Also shown are the optimal (T_opt_, green circles, and adjusted function as green, dashed line) and the maximum (T_max_, triangles) and minimum (T_min,_ inverted triangles) storage temperatures for full achene DR. Increase in DR rate from T_min_ towards T_opt_ is indicated with “□” and decrease in DR rate above T_opt_ towards T_max_ with “□”. Similar notation, in blue, indicates increasing rates of O_2_ uptake and “incomplete” DR at ST>15°C. At MC below ∼4.3% (MC_min_) incomplete DR (white circles) is promoted by ST>35°C. The shaded area indicates tightly bound water forming a monolayer (GAB m_0_ 3.5%). Above the m_0_ and below *T_g_*, multilayer water (second region of isotherms) is in the glassy state; above *T_g_*, multilayer water becomes rubbery. The presence of capillary, free water is inferred from the isotherms when achene MC ∼12-13% and A_w_ ∼0.9 (corresponding to the third region of the isotherm; see **Fig. S 8**). Above this value freezing damage is expected to occur, setting an upper limit for full DR (MC_max_) at sub-zero temperatures. The dotted curve indicates maximum storage temperature that allows a minimum viability of 95% for 90 d (Tmax_(v95%-90d)_) as predicted using the viability equation (Roberts and Ellis, 1980) with sunflower specific constants and Ki 2.32.

The upper MC (MC_max_) threshold for DR is expected to occur where capillary (free) water is present (∼12-13%, within region III of the sorption isotherms, **Fig. 4c**, and **Fig.S8**), and storage at subzero temperatures leads to ice formation and cell damage, or to an active metabolism at ambient temperatures. Within the MC range permissive for full DR, for a fixed MC, increasing ST° between a minimum (ST_min_) and an optimal (ST_opt_) accelerated (and synchronized) DR dynamics at both 10 and 25°C incubation. Supra-optimal ST° further delayed DR (especially at 10°C incubation; **Fig. 5** and Supplementary **Fig. S6-7**). At higher achene MC (∼10%), increasing ST° above 15°C promoted oxygen uptake (**Fig. 4**) together with incomplete DR.

### Delay of DR at supra-optimal ST° vs reduced achene vigor

The bimodal response for full DR observed in the rubbery state (resulting in further delay of DR at supra-optimal ST°) might be explained by a gradual inhibition of reactions involved in this process. Alternatively, an apparent delay of DR could be caused by ageing and lower germination vigor (anticipating loss of viability). To distinguish between these two possibilities (delayed DR vs early loss of vigor), a moisture content shifting experiment was performed (**Figure 8**). The rationale behind this experiment was that if low germination values of achenes stored with high MC (8-10%) at 20°C (which is a supra-optimal ST°) result from deterioration and vigor loss, this would also limit germination after shifting to a lower MC (e.g., 6%). Nevertheless, after drying achenes initially stored with MC 8 or 10% (and which remained dormant after one-month storage) to MC 6%, full DR took place rapidly. This supports that achene vigor was not compromised (i.e., any deterioration was still asymptomatic) during one-month storage of achenes with MC 8-10% at 20°C (supra-optimal ST°), conditions that, otherwise, specifically inhibited DR. On the other hand, increasing MC from 6 to 10% significantly delayed further DR (best observed at 10°C incubation). These different MC levels combined with ST 20°C result in contrasting positions in the state diagram (**Fig. 7**), spanning from the “rubbery” state for MC 6% (optimal for DR) to more fluid states with MC 8 and 10%. Therefore, it can be inferred that reactions leading to DR are favored by a particular rubbery state and inhibited by conditions producing higher mobility.

**Figure 8.**
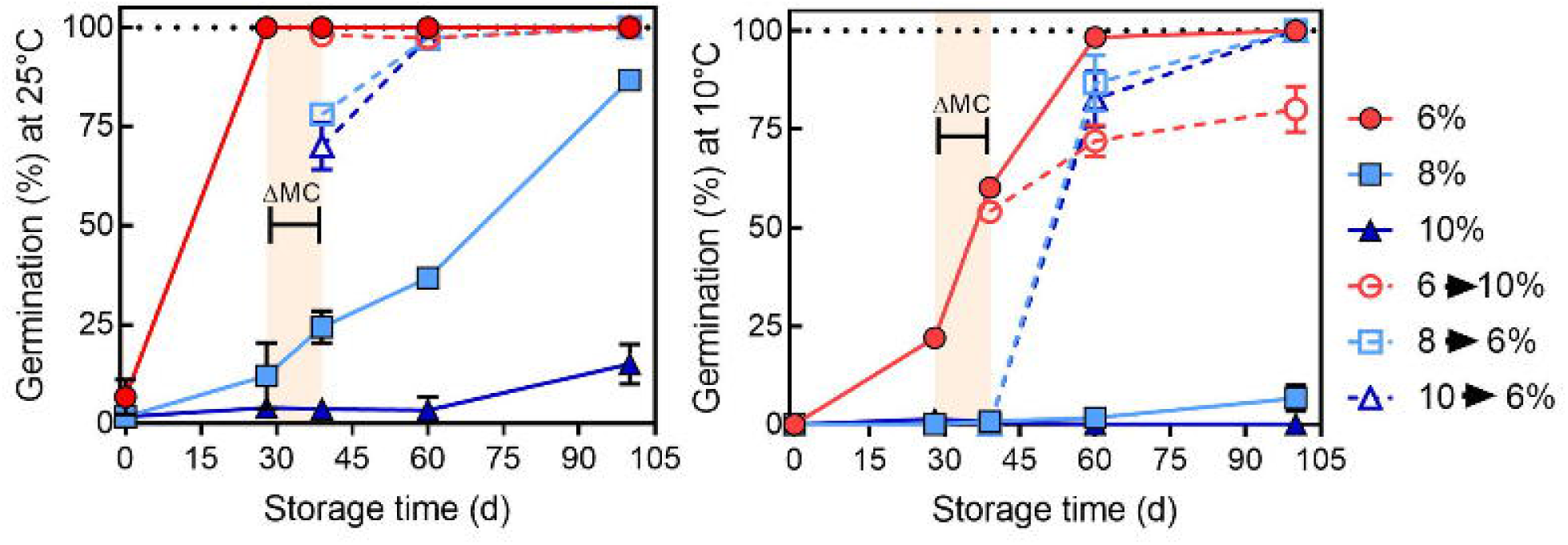
Dormancy release dynamics in moisture content shifting assays. Final germination percent (tested at 25°C and 10°C, left and right panels) after different storage times (0, 28, 38, 60 and 100 days) for achenes with different MC levels which were kept constant (6, 8 or 10% dry weight basis) or were modified after an initial storage of 28 d (empty symbols and dashed lines). On day 28 after storage, achene MC was increased from 6 to 10% (6-+10%; red empty circles) or lowered from 8 and 10 to 6% (8-+6%, squares, 10-+6%, triangles). Increasing MC from 6 to 10% required 11 d in a humid chamber (shaded region in each panel) while drying to MC 6% took only a few hours under air flow at 20°C. Temperature was 20°C during storage and MC manipulation. Each data point is the average of 3 replicate storage containers tested separately. Bars are the S.E.M.

Another MC shifting experiment was performed using dormant achenes with MC 4% stored for 16 months at either 10 or 15°C, conditions that maintained a glassy state and inhibited DR. Nevertheless, after increasing MC to 6% while maintaining each ST° (10 and 15°C), full DR took place within a 40-d period (**Table 1**).

**Table 1:** Effect of increasing MC of achenes after prolonged storage at 10 and 15°C with MC 4%. Achenes with MC of 4.2% (dry weight basis) were first stored for 16 months at 10 or 15°C and presented absolute dormancy. Sub-samples from each pool were humidified to 6%, and continued storage for 40 d at ST 10 or 15°C (as previous). After the second storage period (16 m + 40 d) achene germination was tested at 10 and 25°C. Each value indicates the mean ± S.E.M. (n=3).

| ST<br>(°C) | MC<br>(%) | Incubation<br>T (°C) | Germination<br>(%) |
| --- | --- | --- | --- |
| 10 | 4 | 10 | 0 ± 0 |
|  |  | 25 | 0 ± 0 |
|  | 4→6 | 10 | 97 ± 3 |
|  |  | 25 | 100 ± 0 |
| 15 | 4 | 10 | 0 ± 0 |
|  |  | 25 | 0 ± 0 |
|  | 4→6 | 10 | 100 ± 0 |
|  |  | 25 | 100 ± 0 |

### A two-dimensional MCxST° map for DR in sunflower

The complex patterns observed for DR along this work are summarized in **Fig. 9**. While the glassy-rubbery transition is described by the *Tg*, a threshold between rubbery to fluid is inferred from the physiological data (increase in oxygen uptake, full DR inhibition, and the slope of the longevity line). Within the glassy, rubbery and fluid states, DR is limited to three regions (I, II, III). These are also distinguished according to how DR responds to ST° (positive vs bimodal response) and by the occurrence of full vs incomplete DR. In region I, within the glassy state, DR proceeds in an incomplete way (i.e., achenes can germinate only at warm temperatures, e.g., 25°C) and is promoted by high ST° (between *ca* 35-50°C). In region II (within the rubbery state) full DR is allowed and strong MCxST° interactions are evident, with an optimal ST° decreasing from about 30°C to *ca*.- 18°C as MC increases, and a bimodal response to ST° (DR is promoted by warmer ST° towards an optimal and then delayed at supra-optimal ST°). In region III (under a more fluid state, compatible with partial resumption of metabolism) incomplete DR is promoted by increasing ST° above 15°C. Storage conditions explored in the MC shifting experiments (**Fig. 8** and **Table 1**) are also presented in the map and support the association between DR promotion or inhibition and the physical state of water.

**Figure 9.**
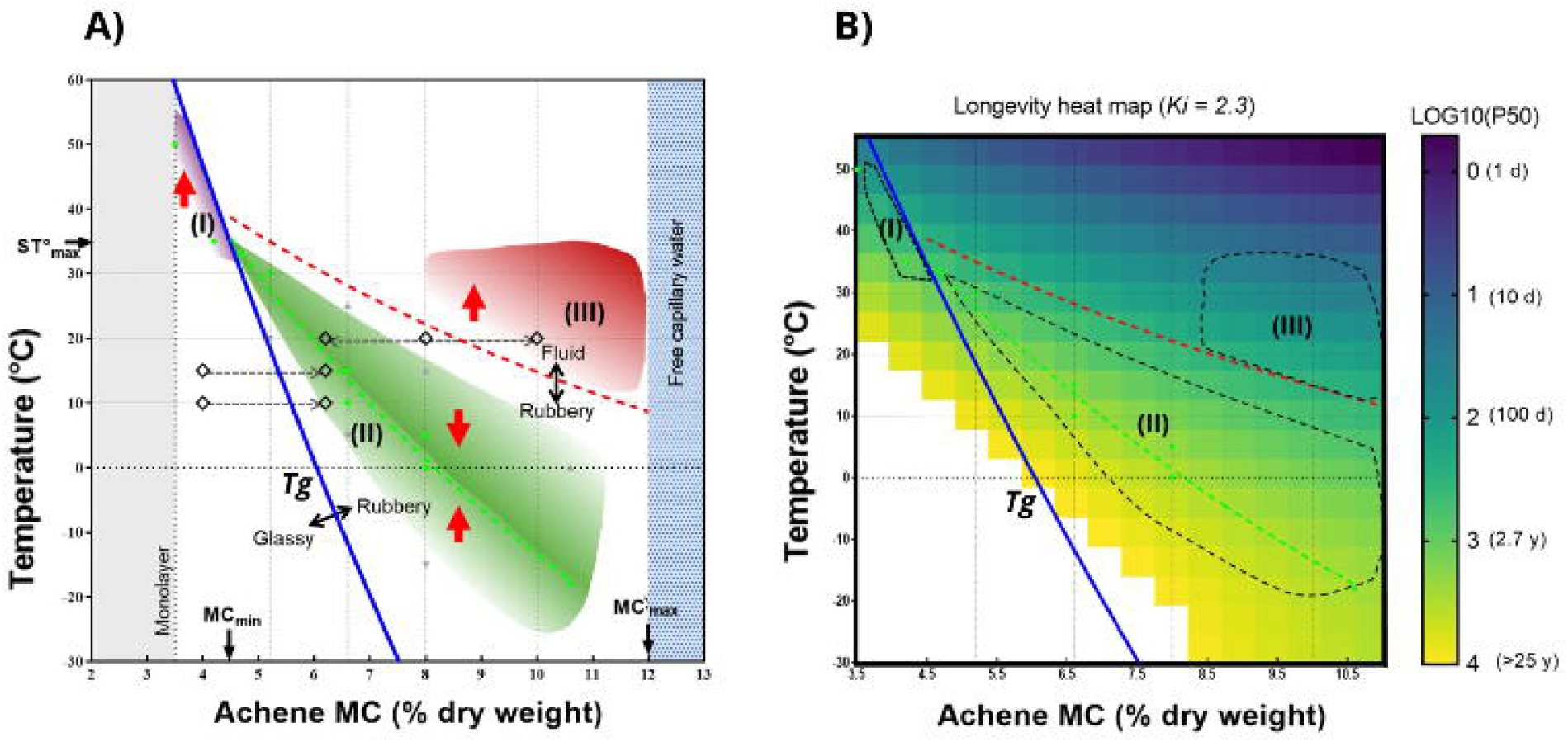
Mapping achene DR and longevity along MCxST° coordinates. **A)** Regions allowing DR (I, II, III) appear shaded in purple (I), green (II) and red (III). Color intensity within each region relates to higher DR rates (either for incomplete DR in regions I and III, or full DR, in region II). Red arrows indicate a positive (upwards) or negative (downwards) response of DR to ST°. Diamonds connected with horizontal dashed lines with arrowheads indicate MC shifting experiments (Fig. 8 and Table 1). Maximum storage temperature (ST_max_), minimum MC (MC_min_) and maximum MC (MC_max_) for full DR in region II are pointed in the MC axis. MC below monolayer (*ca* 3.5%) is shaded in grey. The blue solid line is the *Tg*, and the red dashed line represents a possible threshold between rubbery and fluid (still structured, not capillary) states. **B)** Regions I-III for DR plotted over a predicted longevity heat map. Longevity is expressed as LOG10(P50), where P50 is the time in days to lose 50% viability. Solid green datapoints in A) and B) are the observed optimal ST° for DR along the MC range explored in this work (3.5-10.6%).

## Discussion

In this work we performed a thorough characterization of dormancy release in sunflower achenes under a wide range of MCxST° conditions. A key aspect of our approach was shortening the time to reach different target MC, so that any changes in dormancy level during storage could be attributed to time spent under a fixed MC-ST° condition. Dormancy status was followed over several months, by incubating achenes at 10 and 25°C, as well as embryos in water and ABA. This allowed us to observe that, under some storage conditions, dormancy alleviation proceeded in an incomplete way and achenes could germinate only at warm, e.g., 25°C, but not at cool temperatures, e.g., 10°C. This contrasted with “full” DR, which allowed achene germination along the thermal range (10-30°C). Novel and robust interactions between achene MC and ST° on DR were detected repeatedly in three experimental years. The seemingly complex patterns for DR reported here (Fig. 1) can be explained by how MC and ST° affect the physical state of water in the seed. When the optimal ST° for DR was plotted as a function of achene MC (Fig. 7), a sharp negative association became evident (i.e., ST° optimizing full DR decreased from +35°C to-18°C as MC increased from ∼4.5 to 10%) resembling a state diagram as expected for the water glass transition. *Tg* is the temperature above which water transitions from glassy (amorphous solid) to a much less viscous, rubbery state. *Tg* was measured in embryo axes using two different methods, DSC and ^1^H-NMR, with similar results. The state diagram we obtained is comparable to those reported for embryo axes in maize (Williams and Leopold, 1989), soybean (Bruni and Leopold, 1992) and *Pinus densiflora* (Gerna *et al*., 2022), where the *Tg* falls from over +40°C to below-20°C as MC increases. In contrast, our state diagram differed from the one obtained for sunflower cotyledons by Lehner *et al*. (2006), where *Tg* ranged from 70–80°C in the driest samples to approximately 30°C at 0.14 g H2O g DW^-1^, reflecting large differences in composition between embryo axes and cotyledon cells.

After mapping relevant physiological processes (full and incomplete DR, onset of respiration and viability loss) along MC x ST° coordinates together with the *Tg* (Fig. 9), we propose that DR can proceed, although differently, during storage conditions producing glassy, rubbery and fluid states. According to the model proposed based on our data, reactions leading to full DR are only allowed in the rubbery state (above *T_g_*) and become limited in the glassy but also under more fluid states, where only incomplete DR occurs.

Whether DR is exclusively triggered by oxidative reactions remains unclear. It is widely accepted that oxidative reactions have a key role in the alleviation of seed dormancy during dry storage (Oracz *et al*., 2007; Buijs *et al*., 2018; Morscher *et al*., 2015; C. Bailly, 2019). Nevertheless, in the rubbery state, still in the absence of integrated metabolism, some enzyme activity may be possible, as demonstrated by Candotto Carniel *et al*. (2020) after tracking changes in pigments of the xanthophyll cycle in a lichen. Some degree of enzyme activity in our study is suggested by the increase in ABA content at higher MC and warmer ST°, which may result from the enzymatic hydrolysis of ABA-GE conjugates by β-glucosidases (Xu *et al*., 2014) rather than multistep *de novo* ABA synthesis. The fact that DR proceeded fully under anoxia (and at a similar rate as normoxia) suggests that oxidative reactions may not be essential for DR. Similarly, dormant Arabidopsis seeds after-ripened normally even under storage at 20 MPa gaseous N2 (Buijs et al., 2018). Alternatively, the exposure and sensitivity of dormancy-related targets to oxidative reactions might be enhanced in the rubbery state as compared to the glassy and fluid states. This possibility is supported by work from Bazin *et al*. (2011b) who observed faster DR and higher oxidation levels of mRNA in sunflower seeds stored at 20°C equilibrated with an RH 50% (conditions expected to produce a rubbery state) as compared to seeds stored at RH 33 and 75% (producing glassy and more fluid states, respectively). Under these conditions, even after flushing with N_2_, residual oxygen or ROS in the seed may still be sufficient to promote DR as proposed by Buijs et al. (2018). Further studies are needed to solve the question regarding an essential and exclusive role of ROS in DR.

Our study is not the first to investigate the effects of MC and ST° on DR in sunflower. In a previous study, Bazin et al. (2011a) concluded that the optimal MC for DR increased with temperatures from 15 to 30°C (i.e., optimal ST° for DR increased with MC). When compared to our model, their pattern roughly applies to conditions ranging from the “upper” rubbery (in the supra-optimal ST° range) to a more fluid state. Work by Bazin et al. was performed by testing germination of sunflower naked seeds, not achenes. Also, the different levels of seed MC (ranging from 3 to 12% dry weight basis) were obtained by equilibrating over saturated salt solutions, where gradual changes in MC may have overlapped with changes in dormancy status. Comparison of MC values between both works should also consider that seeds/embryos have a higher oil content and equilibrate with substantially lower MC values than whole achenes. Bazin et al., (2011a) also developed a predictive thermal-time model for DR which, according to our data, applies to conditions promoting incomplete DR in the fluid state (region III).

It is intriguing why reactions leading to full DR are enhanced under a particular rubbery state and become inhibited with increasing fluidity caused either by increasing temperature above an optimal, or by increasing MC. The role of temperature on the velocity of a given reaction or process can be interpreted from the Arrhenius plots, as presented for DR and ageing (**Fig. S10**). While ageing displays an always negative slope (i.e., ageing is always faster with increasing temperature), DR rates are positively and negatively influenced by temperature at both sides of an optimal ST° which depends on MC. The positive slopes in the Arrhenius plot for DR could reflect slower reaction rates involved in DR, or an increased rate for an inhibitory or reversion reaction as temperature increases above an optimal. The value of (T-*Tg*) has been considered as a useful parameter to estimate the kinetics of physical-chemical non-equilibrium changes occurring above *Tg* (M.S. Rahman, 2009). Nevertheless, a constant (T-*Tg*) may not apply to full DR in sunflower, as the adjusted functions for the optimal temperature for full DR and the *Tg* are not parallel. Similarly, Pritchard and Dickie (2003) observed that the *Tg* line is steeper than the mean viability periods in pea and soybean plotted along a MC gradient. These patterns may reflect a more abrupt transition (i.e., a narrow temperature range) from glassy to fluid when MC is low, and a more gradual transition at higher MC. Understanding how properties of the glassy matrix impact on DR requires more detail about molecular relaxations, as provided by mechanical analyses or TDMA (Ballesteros and Walters, 2011; 2019). Ballesteros and Walters (2011) proposed that *Tg* is probably a composite of several types of molecular relaxations and that *Tg* alone does not provide a means to quantify the effects of moisture or temperature on seed deterioration. The TDMA detects changes in visco—elastic properties and can discriminate between large and short-scale movements through the quantification of α and β relaxations, respectively. Future studies using TDMA should provide more detail on the relationship between physical states (allowing different types of molecular motions) and DR. For example, the “rubbery – fluid” transition proposed here could be related to the alpha relaxation, which, as observed in pea cotyledons, runs above the *Tg* and has a smaller slope, both functions converging towards lower MC.

The existence of DR reactions specifically enhanced in the rubbery state (and different from those involved in ageing and DR in glassy and fluid states) is supported by the uncoupling of the observed DR dynamics and the predicted ageing dynamics in this study. Under storage conditions producing a rubbery state, full DR widely anticipated ageing, in contrast to the glassy and more fluid states, where DR (incomplete) was followed more closely by viability loss (**Fig. 9** and **Table S3**). The MC shifting experiment also points at reactions specifically involved in DR and different from those involved in ageing (**Fig. 8**). Further research is needed to understand the nature of these reactions and their targets leading to reduced embryo responsiveness to ABA and dormancy alleviation.

The relationship between DR and the physical state of water reported here may not be restricted to sunflower. Some species are known to dry after-ripen at cold and even sub-zero temperatures (Baskin and Baskin, 2020) and may share a similar mechanism. An inverse relationship between storage temperature promoting DR and MC was reported in *Avena fatua* by Foley (1994). Interestingly, both *A.fatua* and sunflower display remarkable embryo dormancy in addition to coat-imposed dormancy (G.M. Simpson, 1965). The interplay between the embryo and the seed/fruit coats on dormancy expression is complex. Low incubation temperatures enhance the expression of embryo dormancy in sunflower, while inhibition of achene germination at high temperatures is strongly imposed by the pericarp (Corbineau *et al*., 1990). In a study performed with several sunflower genotypes by Arata *et al*. (2021), embryo sensitivity to exogenous ABA was several times (5-10) higher at 10°C incubation as compared to 30°C (e.g., embryo germination was 80% in 50 uM ABA at 30°C, and 0% in 5 uM ABA at 10°C). These previous results together with the ones presented here support the idea that efficient attenuation of ABA signaling in the embryo is instrumental to permit achene germination at cool temperatures, and that the reactions involved are only favored under storage conditions producing a particular physical state of the seed material.

From an ecological perspective, DR of dry achenes at mild (e.g., +5 - 30°C) but also at subzero temperatures (e.g.,-18°C as reported here) is crucial to the adaptation of wild sunflower (*H. annuus* L.) to a wide climatic range. As a summer annual native to the central prairies of North America, *H. annuus* achenes usually germinate in spring after cold or (depending on the latitude) freezing winters (where minimum temperatures can fall below-20°C). Dormancy breaking by dry and cold after-ripening is also compatible with higher longevity, allowing sunflower achenes to remain viable in the soil at least for a few years (Alexander, H.M. and Schrag, A.M., 2003). On the contrary, DR by high temperatures may also be ecologically relevant when moist and warm conditions occur, and a competence between germinating or dying is pushed forward. Clearly, dormancy mechanisms in sunflowers (wild and crop) display an enormous plasticity, with significant effects of the maternal (Riveira-Rubin et al., 2021) and post-dispersal environments as presented here. This adds to the diversity of dormancy mechanisms in the Asteraceae family and successful adaptation to all major vegetation zones worldwide, as reviewed by Baskin and Baskin (2023). Finally, results presented here also provide a solid basis for designing reliable post-harvest protocols to produce high quality commercial sunflower seed, by accelerating dormancy release while maximizing seed longevity.

## Supplementary data

Fig. S1 Moisture content adjustment protocols in experiments 1, 2 and 3. Fig. S2 Germination data (achenes and embryos) in experiment 1.

Fig. S3 Temporal changes in achene germination in experiment 1 including 7- month storage.

Fig. S4 Indicators of vigor and deterioration for achenes stored under different MCxST conditions.

Fig. S5 Images of 7-month storage experiment to assess seed vigor and dormancy Fig. S6 Dormancy release dynamics in experiments 1 and 2

Fig. S7 Additional dormancy release dynamics in experiment 3. Fig. S8 Moisture content isotherms for achenes and embryos Fig. S9 Thermal transition analyses

Fig. S10 Arrhenius plots for dormancy and ageing rates.

Table S1 Published works exploring effect of MC and temperature on dry after-ripening.

Table S2 Pre-storage germination data for achene samples in Figure 1 (experiment 2).

Table S3 Dormancy release and viability loss in the glassy, rubbery and fluid states.

## Abbreviations

ST°: Storage temperature
MC: moisture content
DR: dormancy release
DRR: dormancy release rate
DAS: days after storage
ABA: Abscisic acid
DSC: differential scanning calorimetry
^1^H-NMR: nuclear magnetic resonance
*T_g_*: glass transition temperature
RH: relative humidity
ROS: reactive oxygen species

## Acknowledgements

We are grateful to Cristian Escudero, Maxi Rodríguez and Mirta Tinaro for their valuable technical help during the experiments, and to Dr. Rodolfo A. Sánchez for critical reading of the manuscript.

## Competing interests

The Authors declare that there are no existing competing interests.

## Author contributions

Conceptualization: MVR, GJA and DB. Supervision: MVR. Methodology (germination assays): MVR, PVD, MR-R and GJA. Methodology (DSC and NMR analysis): MPB, FM, GR. Investigation: GJA and MVR. Analysis and visualization of data: MVR and GJA. Writing of original draft: MVR and GJA. Writing and final editing of manuscript: MVR. Funding acquisition: MVR and DB.

## Funding

This work was funded by the Universidad de Buenos Aires (research grants UBACYT No.20020170100599BA, 2018–2021), and by the National Scientific and Technological Research Council of Argentina (CONICET; research grant PIP 2015–2018, No. 11220130100669) and by the National Agency for promotion of Science and Technology (ANPCyT, PICT 2018-3546). Funding acquisition: DB and MVR. This work is part of the doctoral thesis of GJA, as recipient of a PhD fellowship from CONICET under direction of MVR.

## Data availability

All plant materials are publicly available as well as the original data upon request to the authors.

## Notes

### Competing Interest Statement

The authors have declared no competing interest.

